# Gut Microbiota Mediates the Protective Effects of Andrographolide Inhibits Inflammation and Nonalcoholic Fatty Liver Disease(NAFLD) in High-Fat Diet induced ApoE(-/-) Mice

**DOI:** 10.1101/2020.01.24.919316

**Authors:** Shuai Shi, Xin-Yu Ji, Jing-Jing Shi, Shu-Qing Shi, Qiu-Lei Jia, Guo-Zhen Yuan, Qiu-Yan Zhang, Yu Dong, Ying-Dong Lu, Han-Ming Cui, Yuan-Hui Hu

**Author notes:** These authors contributed equally to this study (Shuai Shi, Xin-Yu Ji, Jing-Jing Shi). Correspondence authors Yuan-Hui Hu, Guang’an Men Hospital, China Academy of Chinese Medical Sciences, No. 5, North Line Court, Road, Xicheng District, Beijing 100053, People’s Republic of China, Han-Ming Cui, Guang’an Men Hospital, China Academy of Chinese Medical Sciences, No. 5, North Line Court, Road, Xicheng District, Beijing 100053, People’s Republic of China, Ying-Dong Lu Guang’an Men Hospital, China Academy of Chinese Medical Sciences, No. 5, North Line Court, Road, Xicheng District, Beijing 100053, People’s Republic of China.

## Abstract

Mechanisms relating the gut bacteria to Nonalcoholic Fatty Liver Disease (NAFLD) have been proposed containing the dysbiosis-induced dysregulation of hepatic lipid metabolism that allows for the translocation of microbial components and leads to hepatic inflammation and steatosis. Andrographolide (AG) regulates inflammation mediated by NF-κB pathway which also play a key role in reduction of inflammation and fibrosis in experimental nonalcoholic steatohepatitis (NASH), yet the mechanisms linking this effect to gut microbiota remain obscure. Here we show that ApoE knockout (Apoe -/-) mice fed a high-fat diet (HFD) supplemented with AG regulates levels of biochemical index and inflammatory cytokines associated with gut microbe. Moreover, HEPG2 cells induced by ox-LDL were used as validation in vitro. H&E staining and Oil-Red staining were respectively used for tissue and cells morphology. Gut microbiota were examined by 16S rRNA sequencing. Expression of NF-κB, C/EBPβ and PPAR-γ in liver and HEPG2 cells were detected by western blot and qRT-PCR. The results showed, among others, that AG alleviate hepatic steatosis and fat content in HEPG2 cells, while it induced decreased levels of Bacteroides, and increased levels of Faecalibaculum, Akkermansia. We further identified that inhibition of NF-κB/C/EBPβ/PPAR-γ pathway of hepatic steatosis model in vivo and vitro by AG also contributes to prevention of HFD-induced inflammation and dislipidemia. Importantly, as result of pearson correlation, Bacteroides may be the most relevant one fundamentally involved in the mechanism of AG attenuates NAFLD. Together, our findings uncover an interaction between AG and gut microbiota as a novel mechanism for the anti-NAFLD effect of AG acting through prevention of microbial dysbiosis, dislipidemia and inflammation.

**Importance:** HFD due to gut microbial dysbiosis is a major contributor to the pathogenesis of dislipidemia and inflammation, which primarily mediates the development of NAFLD. A treatment strategy to reduce both dislipidemia and inflammation appears to be an effective approach for addressing the issue of NAFLD. Andrographolide (AG) is the major effect component in traditional Chinese medicine Chuan-xin-lian (Andrographis). Little is known about the role of gut microbiota in the anti-NAFLD effect of AG. 16S rRNA gene sequencing revealed that AG significantly decreased Bacteroides and increased Faecalibaculum, Akkermansia. By using vivo and vitro experiment, we prove that gut microbiota plays a key role in AG-induced protective against high-fat-diet-induced dislipidemia and inflammation. Moreover, NF-κB/C/EBPβ/PPAR-γ pathway inhibition was partially involved in the beneficial effect of AG. Together, these data suggest that the gut microbiome is a critical factor for the anti-NAFLD effects of AG.

## Introduction

Nonalcoholic fatty liver disease (NAFLD) is characterized by an increase in the accumulation of lipids in the liver. Steatosis can progress to steatohepatitis and clinically significant fibrosis(1). There is now growing evidence that NAFLD is a multisystem disease. In United States, NAFLD-cirrhosis or NAFLD-Hepatocellular Carcinoma (HCC) are now the second cause of liver transplantation (2). It also has been shown that the clinical burden of NAFLD is not only confined to liver-related morbidity and mortality, but also affected extra-hepatic organs and regulatory pathways such as NAFLD increases risk of type 2 diabetes mellitus (T2DM), cardiovascular (CVD) and cardiac diseases, and chronic kidney disease (CKD). A large cohort study of adults with no history of CVD, NAFLD was significantly associated with the development of coronary artery calcium independent of cardiovascular and metabolic risk factors (3). It is worth noting that the majority of deaths among NAFLD patients are attributable to CVD (4). Like many Western countries, urbanisation in many Asian countries has led to significant changes in lifestyle and eating behaviors in last two decades. Epidemiological studies have showed prevalence of NAFLD in Asia was around 25% (5). The molecular mechanisms of hepatic steatosis in NAFLD, attributing to the four major pathways contributing to lipid homeostasis in the liver (6). Development of NAFLD is determined by gut bacteria. Mechanisms relating the bacteria to NAFLD have been proposed containing the dysbiosis-induced dysregulation of gut endothelial barrier function that allows for the translocation of microbial components and leads to hepatic inflammation (7). Non-drug therapies including reasonable calories diet, micro- and macronutrients and physical activity and exercise are benefit to the treatment of NAFLD (8). Additionally, NAFLD are commonly considered to treating by lipid-lowering drugs like statins (9). Potential adverse effects such as myopathy and hepatotoxicity are associated with statins and its drug-drug interactions (10). Hence, the primary task are still looking for safer and more effective drugs.

For centuries, traditional herbal medicine are used to treat a myriad of maladies, and simultaneously made great achievements (11)(12). Bioactive compounds from herbal medicine participate reduction of cholesterol synthesis, elevation of reverse cholesterol transport, and promotion of cholesterol excretion in the liver (13). Andrographis paniculata has hundreds of years been part of the traditional Chinese medicine in China. Andrographolide(AG) was isolated as a major bioactive ingredient of it in 1951. Since 1984, AG have been evaluated with modern drug discovery approach for anti-inflammatory effects, and it was found to exert cytotoxic/anticancer effects with the underlying miscellaneous actions (14)(15). Treatment with AG dose dependently suppressed cardiac inflammation mediated by NF-κB pathway which also play a causal role in reduction of inflammation and fibrosis in experimental nonalcoholic steatohepatitis (NASH) (16)(17).AG inhibits macrophage foam cell formation caused by two mechanisms which are inhibition of CD36-mediated Ox-LDL uptake and induction of ABCA1- and ABCG1-dependent cholesterol efflux. It could be a potential candidate to prevent atherosclerosis (AS) (18).New type of AG which loaded the block copolymer of poly (ethylene glycol) and poly (propylene sulphide) micelle is considered to synchronically alleviate inflammation and oxidative stress, and provide a promising and innovative strategy against atherosclerosis (19). Sterol regulatory-element binding proteins (SREBPs) control Cellular lipid metabolism and homeostasis. They are implicated in complicated pathogenic processes such as endoplasmic reticulum stress, inflammation and autophagy, and as a result, they contribute to obesity, dyslipidemia, diabetes mellitus, AS and NAFLD (20). AG regulated SREBP and other metabolism-associated genes in liver or brown adipose tissue, which may have directly contributed to the lower lipid levels and enhanced insulin sensitivity in mice with high fat diet (HFD)-induced obesity (21). Decrease of pathological lesions contain AS and NAFLD in ApoE knockout (ApoE -/-) mice with HFD can be controlled by regulating the lipid metabolism and inflammatory (22). We tried to intervene it by AG to analyze the information from gut microbes.

## Materials and methods

### Animal model and study design

Six-week-old male C57Bl/6J (WT group, n=8) and ApoE−/− mice(knockout from C57Bl/6J, n=40) were provided by Beijing Vital River Laboratory Animal Technology Co., Ltd (Beijing, China). All the mice were housed in microisolator cages and assayed under conditions of controlled temperature of 24±2 °C and humidity of 60% in a 12-h light/dark cycle with free access to standard chow and sterile water.

After 1 week of acclimatization to the environment, 40 male ApoE−/− mice were randomly divided into 5 groups: High Fat Diet group (HFD group, n = 8), high fat diet plus atorvastatin calcium (LOT. X80827, Pfizer, Dalian, China) group (ATO group, n =8), high fat diet plus high dose AG group (AGH group, 90mg/kg/d, n=8), high fat diet plus middle dose AG group (AGM group, 30mg/kg/d, n=8) and high fat diet plus low dose AG group (AGL group, 10mg/kg/d, n=8). AG was extracted as high purity crystalline powder with inspection Report (No. C-01-19013) by a drug company (LOT. 190402, Sichuan Wenlong Pharmaceutical Co., Ltd, Sichuan, China).

5 groups received high fat diet (Catalog No. H10141, Beijing HFK Bioscience CO.,LTD) for 8 weeks, and meanwhile, 8 male C57Bl/6J mice as healthy control were fed chow diet from SPF (Beijing) biological technology co,. LTD (Beijing, China). Details of the feeds is reflected in the supplementary table 1.

Body weight and food intake were assessed weekly. Fecal pellets were collected at 4 weeks and 8 weeks from each mouse directly into individual sterile microcentrifuge tubes and stored at -80°C for DNA extraction and analysis. 8 weeks later, All the mice were anesthetized (Serial No. V942265, Matrx VIP 3000 Isoflurane Vaporizer, New York, USA) after overnight fasting. Retro-orbital blood samples were collected in Vacuette Blood Collection Tubes (REF:454078CN, Greiner Bio-One, Thailand) for serum. Part serum from every mice were immediately detected for biochemical analysis and others were storage at -80°C for inflammatory cytokines. Liver tissues were immediately excised under sterile conditions followed by weight determination and washing with ice-cold normal saline, and then fixed in 4% PFA for pathological detection.

### Biochemical analysis

Blood samples were collected from all the mice. Serum was centrifuged at 12000 g and 4 °C for 15 min (23). To avoid the impact of the freezing, serum total cholesterol (TC), triglycerides (TG), high-density lipoprotein cholesterol (HDL-C), low-density lipoprotein cholesterol (LDL-C), aspartate aminotransferase (AST), alanine aminotransferase(ALT), creatinine (Cr) and blood urea nitrogen (BUN) was immediately measured by Beckman Coulter series chemistry analyzers (AU5822, Beckman, USA). The assay kits were purchased from Beckman Coulter as supplementary table 2.

### Determination of serum inflammatory cytokines

Frozen serum were thawed and centrifuged for 10 minutes at 10,000 rpm. Meanwhile, warm up the Bio-Plex System at least 30min. Reconstitute a single vial of standards in 500 μL of a diluent similar to the final sample type or matrix. Vortex for 5 sec and incubate on ice for 30 min. Prepare a fourfold standard dilution series and blank. Vortex for 5 sec between liquid transfers. Vortex the 10x single beads for 30 sec and dilute to 1x in Bio-Plex Assay Buffer. Protect from light. Then ran the assay in accordance with the specification (Bio-Plex Pro Mouse Cytokine Grp, Catlog NO. #M60009RDPD, LOT. 64212941, Bio-Rad Laboratories, Inc., USA). Read the plate at Bio-Plex MAGPIX System (Bio-Rad Laboratories, Inc., USA). Finally, calculated interleukin-1α (IL-1α), interleukin-1β(IL-1β), interleukin-6 (IL-6) and tumor necrosis factor α (TNFα) concentration by the standard curve.

### Histology of liver

The liver tissue samples were 4% PFA fixed and embedded in paraffin, and 8 µm-thick paraffin sections were stained with the hematoxylin & eosin(H&E) method as described previously (24). Finally, tissue was observed by a IX70 light microscope (Olympus, Tokyo, Japan) equipped with an imaging system at 200 magnification.

### Gut microbiota composition

Gut microbiota analysis was sequenced by construct a library of small fragments using the method of Paired-End sequencing based on the Illumina HiSeq sequencing platform.Composition of species was revealed by splicing and filtering the Reads, Operational Taxonomic Units (OTUs) cluster, species annotation and abundance analysis. Furthermore, Difference between groups can be analyzed through Alpha Diversity, Beta Diversity and species difference analysis, etc. Genomic DNA was extracted from caecal content using a PowerSoil^®^ DNA Isolation Kit (Catalog No. 12888-100, MO BIO Laboratories, Inc., USA). The V3-V4 region of the 16S rDNA gene was amplified by PCR with modified primers sets for the Illumina HiSeq 2500 sequencing instrument. The PCR was performed in a total reaction volume of 20 µL as described (25). Paired-end reads were merged by FLASH (26) ^(^version 1.2.11). Raw Tags were filtered by Trimmomatic (27), (version 0.33) for clean tags. Chimeric sequences were identified and removed through UCHIME (28) (version 4.2). Effective Tags were finally obtained and clustered into OTUs at a 97% similarity threshold (USEARCH, version 10.0)

OUT were filtered at 0.005% in all sequences (29).The library construction, sequencing and QIIME (30) analysis (version 1.8.0) were performed by Beijing Biomarker Technologies Co., Ltd., (Beijing, China).

#### Cell culture and treatment

The human hepatoma cell line HepG2 was obtained from the Cell Resource Center, Peking Union Medical College (which is the headquarter of National Infrastructure of Cell Line Resource, NSTI) on Mar.15th, 2018. The cell line was checked free of mycoplasma contamination by PCR and culture. Its species origin was confirmed with PCR. The identity of the cell line was authenticated with STR profiling (FBI, CODIS). All the results can be viewed on the website (http://cellresource.cn).

HepG2 cells were grown in a CO_2_ incubator (5% CO_2_, 95% air) in a 25-cm^2^ cell culture flask with 2μm vent cap (Cat No. 430639, Corning Incorporated, China). Cultures were maintained in 20 mL MEM-NEAA (Cat No. CM50011, Lot No. D2215522, CELL technologies, China) containing 10% FETAL BOVINE SERUM DEFINED (FBS, Cat No. SH30070.03, Lot No. AYC60564, Hyclone laboratories., lnc, USA), 1.25%L-glutamine, and 2% penicil-lin/streptomycin. About 5 or 6 days before each experiment, 1million cells were seeded in 6-well tissue clusters (Costar) in 3 mL DMEM containing 10% FBS (31). Ox-LDLs were obtained from Shanghai yuanye Bio-Technology Co., Ltd (Cat No. S24879-2mg, Lot No. L29O9W73395, China). Cells in 6-well tissue clusters were divided into 6 groups: untreated cells (Contrast group, CON), ox-LDL treat cells (Model group, MOD), ox-LDL+ 10μmol/L atorvastatin calcium (Cat No. 134523-03-8, Lot No. J0303AS, Dalian Mellunbio. Co., Ltd. Manufacture, China) treat cells (ATO group), ox-LDL+ high dose (50μmol/L) AG treat cells (AGH group), ox-LDL+ middle dose (5μmol/L) AG treat cells (AGM group), ox-LDL+ low dose (0.5μmol/L) AG treat cells (AGL group). Cells at 80% confluence were stimulated with 50μg/mL ox-LDL for 24 h. The final concentration of ox-LDL was decided by preliminary experiments and previous research (32). After induced by ox-LDL, cells were respectively treated with AG as high dose, middle dose and low dose.

### Oil-Red staining

Oil-Red O powder (Cat No. O0625-100G, Lot No. #SLBM444V, SIGMA-ALDRICH, USA) was prepared according to the description. Cells after treated in the indicated groups were fixed with 4% PARAFORMALDEHYDE (PFA, Ref No. BL539A, Lot No. 69100900, biosharp, Beijing Labgic Technology Co., Ltd, China) for 30 min and then stained by oil-red O for 15 min. Finally, cells was observed by a IX70 light microscope (Olympus, Tokyo, Japan) equipped with an imaging system at 200 magnification.

### SDS-PAGE and western blot

The methods of protein extraction from the liver tissues and cells, electrophoresis, and subsequent blotting were conducted as previously described with some modifications. The homogenized tissues were lysed with RIPA buffer (high) (Cat No. #R0010, Lot No. 20190318, Beijing Solarbio Science & Technology Co., Ltd, China) containing halt protease and phosphatase inhibitor single-use cocktail for 30 min on ice. The lysates were centrifuged at 12,000 revolution for 30 min at 4°C. Each sample protein was separated on TGX Stain-Free FastCast Acrylamide Kit, 10% (Cat No. #1610183, Batch No. 64278801, Bio-Rad Laboratories, Inc., www.bio-rad.com, USA) and immunoblotting with antibodies (Supplementary Table S4.) followed by HRP-conjugated secondary antibody. Visualized bands were analyzed on ChemiDoc™MP imaging system (Model No. Universal Hood III, Serial No. 731BR02915, Bio-Rad Laboratories, Inc., www.bio-rad.com, USA) using ImageLab™software. Quantification was performed with Image Pro Plus software.

### Real-time quantitative polymerase chain reaction

The levels of NF-κB-p65, C/EBPβ and PPARγ mRNA transcripts to control GAPDH were determined by RT-qPCR. Total RNA was extracted from the frozen mice liver tissues with TRIzol Reagent (Ref No. 15596026, Lot No. 190906, ambion by life technologies, USA) according to the manufacturer’s instructions from Direct-zolTM RNA MiniPrep (Cat No. R2052, Lot No. ZRC000661, www.zymoresearch.com, USA). Amount of total RNA was detected by NANODROP 2000 Spectrophotometer (ND2000, Thermo Scientific, Gene Company Limi ed,). cDNA synthesis was performed using the High Capacity cDNA Reverse Transcription Kit (Ref No. 4368814, Lot No. 00643655, appliedbiosystems by Thermo Fisher Scientific, Thermo Fisher Scientific Baltics UAB, Lithuania). Real-time qPCR was performed with the ABI 7900 system (applied biosystems, USA) using an Power SYBR Green PCR Master Mix (Ref No. 4367659, Lot No. 1804574, applied biosystems by Thermo Fisher Scientific, Life Technologies LTD, UK). The primers were synthesized by Thermo Fisher and the sequences were listed in Supplementary Table S5. The thermal conditions of PCR were as follows: 1 cycle of 95°Cfor 10 min, 45 cycles of 95°C for 15 s, 55-60°C for 20 s, and 72°C for 30 s. The relative expression level of each gene was determined by the 2^−ΔΔ^Ct method and normalized to the GAPDH mRNA in each sample (33).

### Statistical analysis

Statistical analysis was performed using SPSS software 13.0 (IBM, Almon, NY, USA). One-way analysis of variance (One-way ANOVA) was performed when multiple group comparison was carried out. Results were expressed as mean ± SD. Results were classified into three significant levels using the *p* value of 0.05, 0.01 and 0.001. Results were classified into one significance level using the *p* value of 0.05. Graphics were presented using GraphPad Prism 8.3.

## Results

### Effect of AG on body weight, food intake and liver weight

Three dosages of AG were administered orally to different groups of high fat diet-induced NAFLD for 8 weeks, and the impact on body weight, food intake and liver weight was determined. The initial mice body weight and the food intake were basically same among all groups. 8 weeks later, body weight in HFD group was significant higher than the other groups (Fig. 2A, p<0.001). Food intake also increased, and the highest growth rates were observed in the WT group (Fig. 2B, p<0.001). Compared with HFD group, food intake in AGH and AGM groups were significant lower in week 8 (Fig. 2B, HFD vs AGH: p<0.01, HFD vs AGM: p<0.05). As shown in Fig. 2C, liver weight in HFD group displayed significant changes among ATO and AGH groups (p<0.05, respectively).

**Figure 1.**
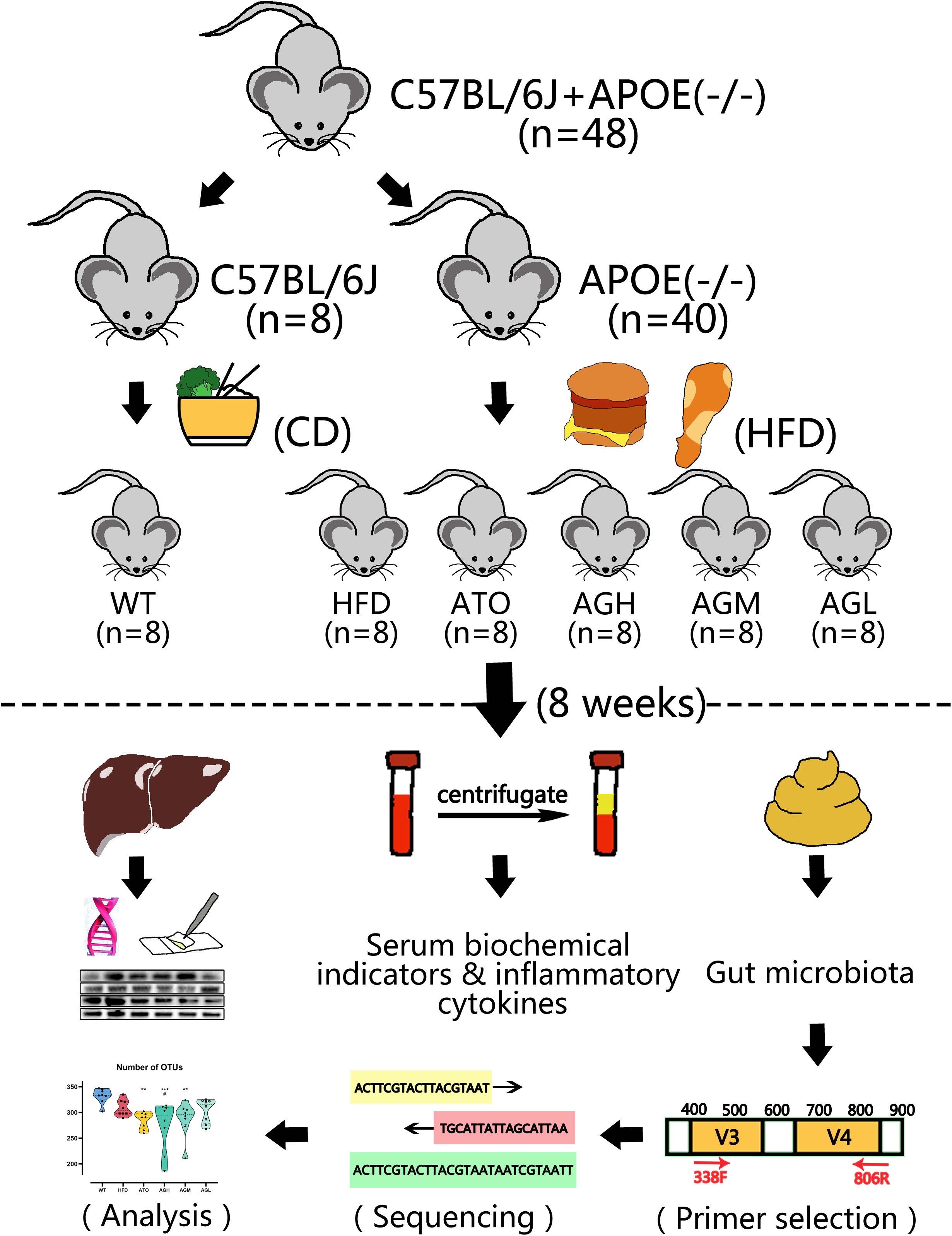
Study design of animal experiment.

**Figure2.**
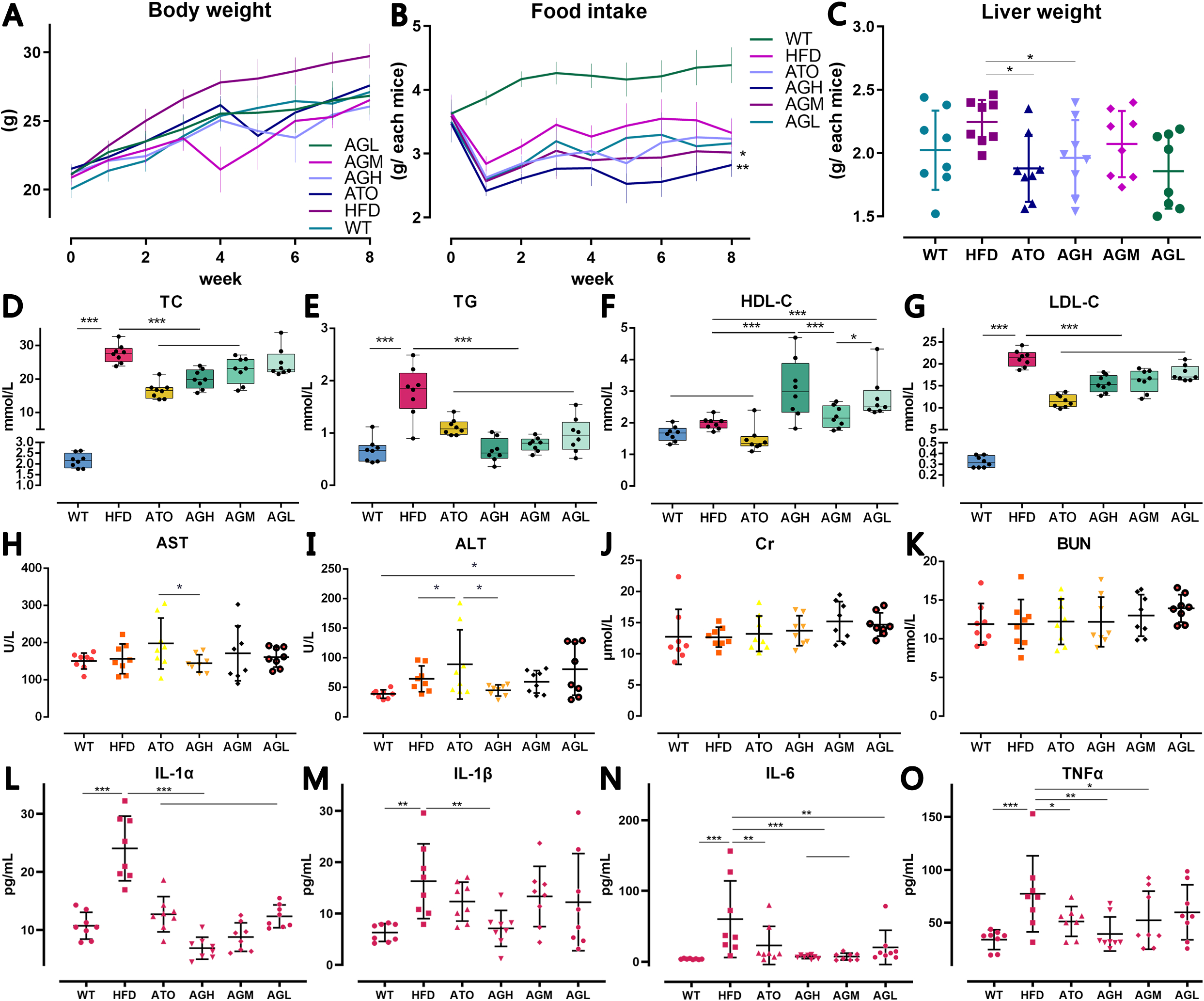
Effect of AG on physical data, serum biochemical indicators and inflammatory cyt okines. **(A)** Body weight. HFD group showed significant higher than other groups at week 8 (p<0.001); **(B)** Food intake in 24h. WT group was significantly fed more among other groups (p<0.001). Mice in AGH and AGM group showed less food intake than HFD group at week 8(AGH: p<0.01, AGM: p<0.05); **(C)** Liver weight; **(D)** Level of total cholesterol (T C); **(E)** Level of triglycerides (TG); **(F)** Level of high-density lipoprotein cholesterol (HDL-C); **(G)** Level of low-density lipoprotein cholesterol (LDL-C); **(H)** Level of aspartate aminotran sferase (AST); **(I)** Level of alanine aminotransferase (ALT); **(J)** Level of creatinine (Cr); **(K)** Level of blood urea nitrogen (BUN); **(L)** Level of interleukin-1α (IL-1α); **(M)** Level of interl eukin-1β (IL-1β); **(N)** Level of interleukin-6 (IL-6); **(O)** Level of tumor necrosis factor α (T NFα). The data are expressed as the mean±SD of the mean (n=8 per group) and analyse d by One-way ANOVA: *p<0.05, **p<0.01, *** p<0.001. C57BL/6J mice were fed a chow diet (WT), ApoE(-/-) mice were fed a high-fat diet (HFD), a high-fat diet and atorvastati n calcium (ATO), a high-fat diet and a high dose of AG (AGH), a high-fat diet and a mid dle dose of AG (AGM) or a high-fat diet and a low dose of AG (AGL) for 8 weeks.

### Effect of AG on the biochemical indicators

As shown in Fig. 2D-G, the serum TC, TG and LDL-C levels in the WT group were much lower than those in the HFD group (p<0.001). The serum HDL-C level exhibited the opposite trend to the LDL-C levels (p>0.05), which were significantly increased in AGH, AGL groups (p<0.001, compared with HFD group). However, no difference were observed among the WT, HFD and ATO groups, may be related to significant changes in total cholesterol in HFD and ATO groups. Serum AST and ALT levels were also detected. Compared with the AGH group, ATO had higher AST and ALT levels with significant difference (Fig. 2H-I, p<0.05). Moreover in ALT, statistical comparisons between HFD vs ATO and WT vs AGL were observed significant difference(Fig. 2I, p<0.05). No significant difference in Cr and BUN was observed (Fig. 2J-K, p>0.05).

### Effect of AG on serum inflammatory cytokines

IL-1α, IL-1β, IL-6 and TNF-α concentration in serum are showed in Fig. 2L-O. IL-1α, IL-1β and TNFα in HFD group were significant higher than WT group (p<0.001). Both atorvastatin and all dose of AG inhibit the increase of IL-1α(Fig. 2L, p<0.001). As shown in Fig. 2M, only AGH group was observed reduce effect with significant difference (p<0.01). Similar to the previous trend, ATO and AGL group significantly decreased serum IL-6 level (Fig. 2N, p<0.01) compared with HFD. AGH and AGM also obviously inhibited the expression of IL-6, more significant difference was observed (p< 0.001). In addition, the expression of TNF-α was reduced by ATO, AGH and AGM with significant difference (Fig. 2O, ATO vs HFD: p<0.05; AGH vs HFD: p<0.01; AGM vs HFD: p<0.05). As a result, both ATO and AG might suppress the relate inflammatory cytokines in the serum of the ApoE-/- mice.

### Evaluation of liver histology

H&E staining of samples from the HFD group showed severe liver steatosis and were characterized by diffusely mixed sizes of fat bubble, especially tend to have big fat bubbles. Meanwhile, H&E staining of samples from the WT group showed almost none fatty changes. After 8 weeks of AG intervention, the fatty proportion of the liver was significantly dose-dependent reduced in the AGL, AGM, AGH and ATO groups, but occasionally showed scattered lipid drops. It is showed a distinct increase in fat accumulation in the liver tissue induced by high fat diet, whereas AG decreased the fat deposition in the liver sections (Fig. 3).

**Figure3.**
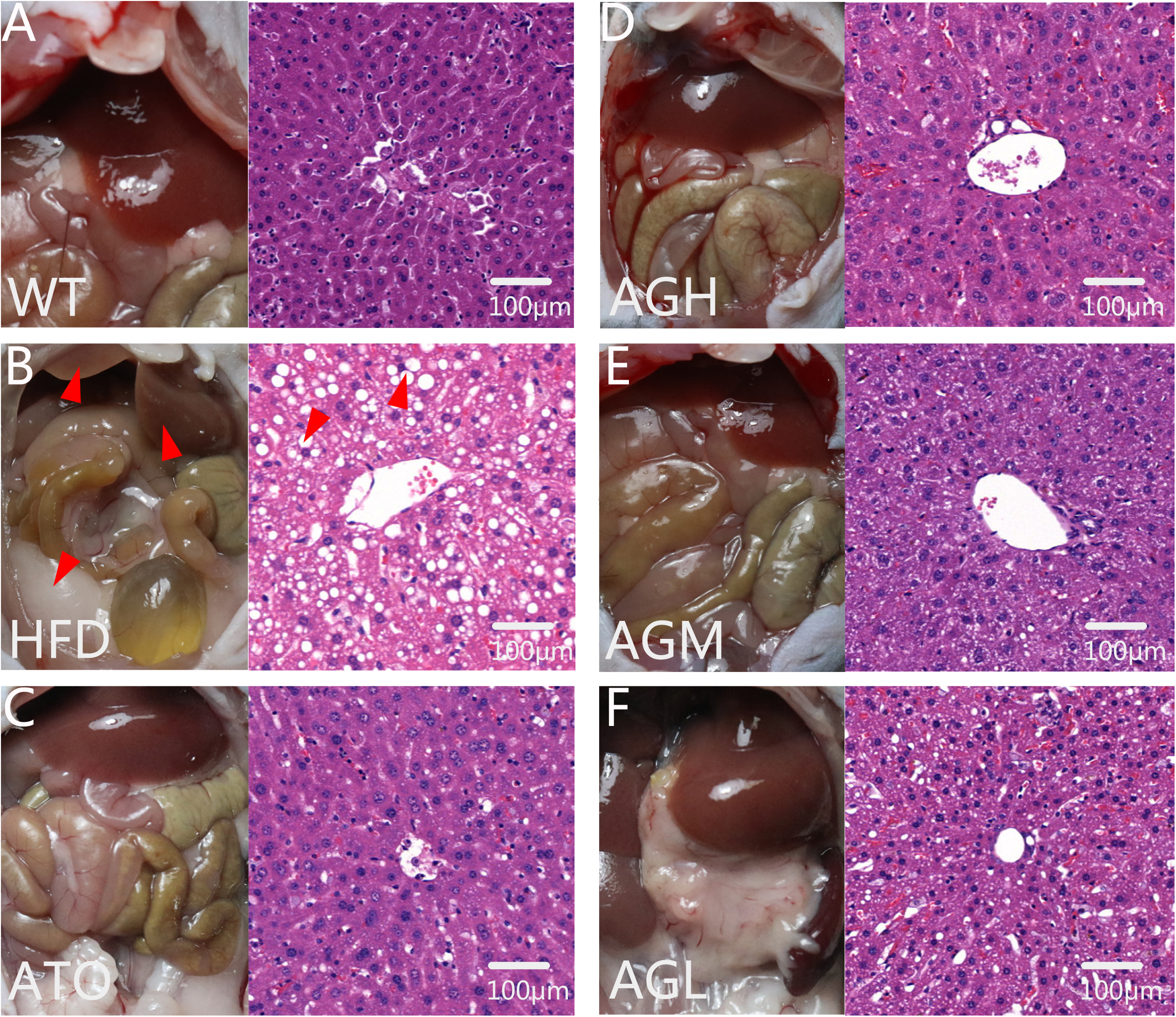
H&E staining of liver sections in all groups (200× magnification). **(A)** H&E staining of WT group; **(B)** H&E staining of HFD group; **(C)** H&E staining of ATO group; **(D)** H&E staining of AGH group; **(E)** H&E staining of AGM group; **(F)** H&E staining of AGL group.

### Changes in the gut microbiotas

The HFD group in previous results showed obviously enhances lipid deposition effect, and mice in ATO, AGH, AGM and AGL were able to know varying degrees of treatment effect. We further explore the gut microenvironment for trying to explain the phenomenon. To assess the change among groups on gut microbiota, we sequenced V3+V4 amplicons of 16S rRNA genes. A total of 3,357,231 high-quality reads were obtained from 3,839,233 reads in 48 samples, with an average of 69,942 reads (at least 52,337) per sample. After comparing the results with the reference database, we performed bacterial taxonomic identification. Then, based on the results of bacterial taxonomic identification, we conducted the following analyses: alpha diversity, beta diversity, visualization and significance analysis of differences among groups, and screening of taxa with significant differences among groups.

441 OTUs were finally obtained. Mice in ATO, AGH and AGM groups exhibited significant losses of OTUs compared to the number of OTUs in WT group in Fig. 4A (ATO vs HFD: p<0.01; AGH vs HFD: p<0.001; AGM vs HFD: p<0.01). AGH treatment mice had less microbes at week 8 than the HFD group mice (p<0.05). Each OTU corresponding to different bacterial species. Rarefaction curve was consist of number of sequences and OTUs. The gradually smooth curve as Fig. 4B shown represent the amount of data from sequencing is sufficient. Shannon curve reflect the diversity of microorganisms in samples, and also prove the amount of data. It can be learned by Fig. 4C that diversity of microbe in ATO, AGH and AGM groups is obviously reduced. In addition to abundance of gut microbiota, evenness can also evaluated by Rank-abundance curve. Fig. 4D was showed compared with other groups, there was more microbial species in WT group which is also more evenly distributed. The OTU-based principal component analysis (PCA) observed the AG-related change is mostly represented by PC1, which explained 75.52% of the variation in the microbial composition; the diet-related difference is represented by PC2, which explained an additional 6.39% of the total variation (Fig.5H). The difference in gut microbiota were not revealed community-wide but observed particular groups of microbes. According to abundance of genus level as Fig.5A-G shown, Bacteroides, Faecalibaculum and Akkermansia may be fundamentally involved in the mechanism of AG attenuates NAFLD. Pearson correlation coefficient showed the efficiency of lipid-lowering and anti-inflammation is mostly correlated to Bacteroides, which is the central gut germ in regulation of AG (Fig.5I).

**Figure4.**
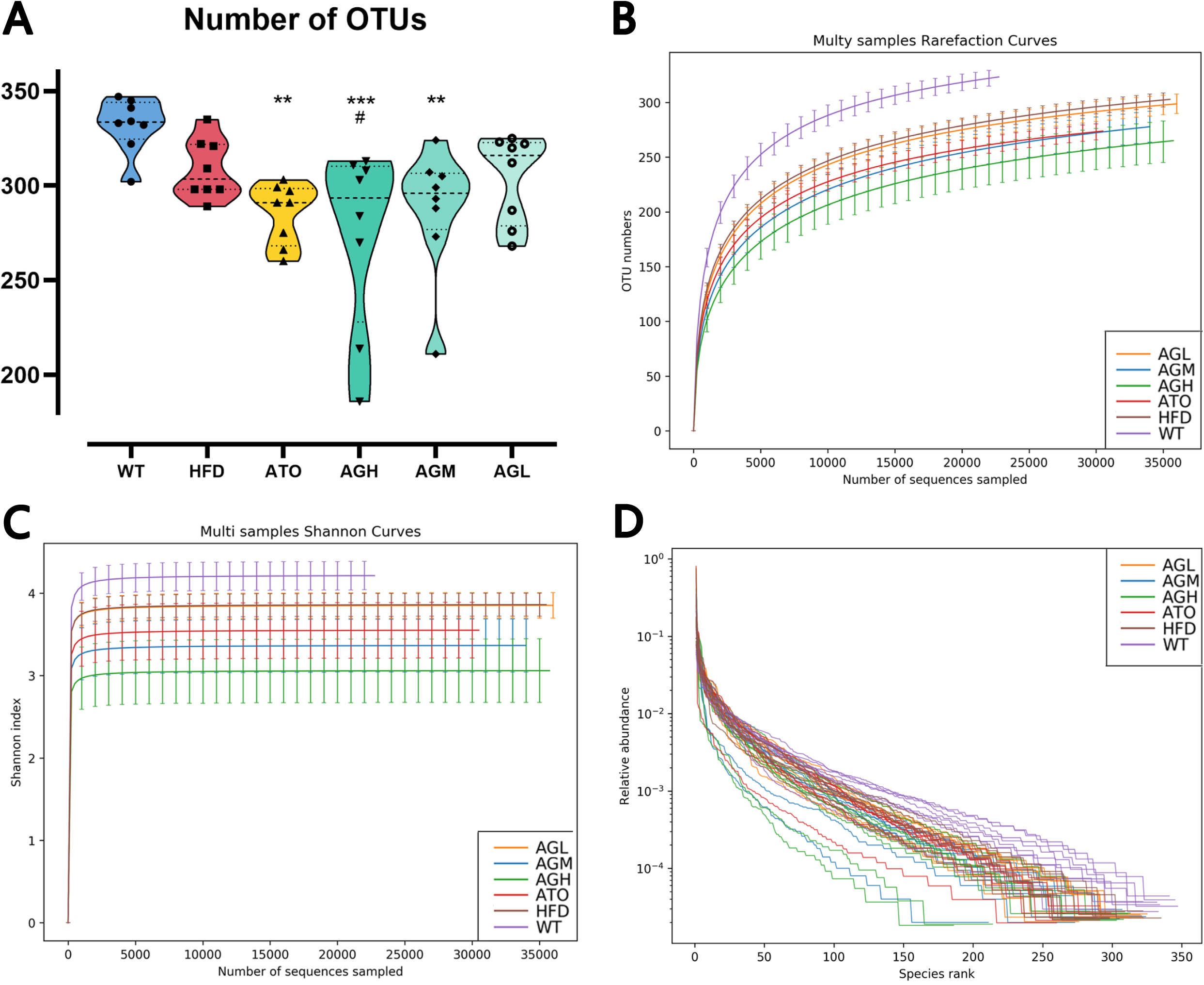
Alpha Diversity of gut microbiota. **(A)** Number of OTUs; **(B)** Rarefaction Curves; **(C)** Shannon-Wiener Curves; **(D)** Rank-Abundance Curves.

**Figure5.**
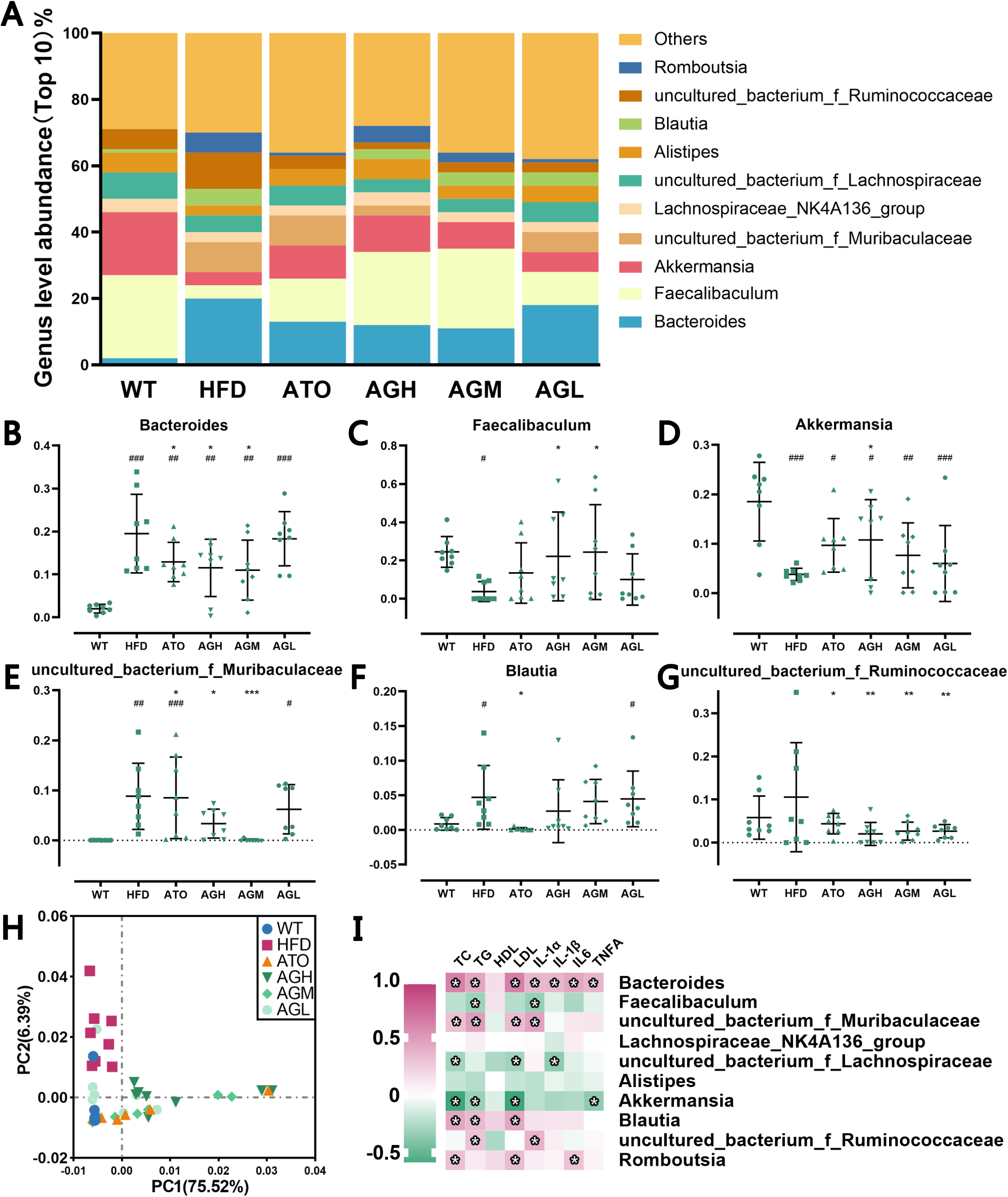
Relative abundance of microorganisms in genus level, Beta Diversity and heatmap of correlation. **(A)** Median relative abundance of microbes in genus level. **(B-G)** Comparison of gut microbiota among 6 groups of mice. T-test was applied to detect the statistical significance. The data are expressed as the mean ± SD of the mean (n=8) and analysed by One-way ANOVA: *p<0.05, **p<0.01, *** p<0.001 versus HFD group, #p<0.05, ##p<0.01, ### p<0.001 versus WT group. **(H)** Principal component analysis (PCA) for gut microbiota of four groups of mice. **(I)** Heatmap representation of the Pearson’s r correlation coefficient between serum lipid, inflammatory cytokines and bacterial taxa in genus level. Only the bacteria, for which at least one significant correlation to serum indicators was found, are displayed. *Adjusted p value <0.05.

### Effect of AG on mRNA and protein expression in liver and HEPG2 cells

To assess the mechanism of AG for reducing the hepatic inflammation and lipid deposition, the protein expression and transcription levels of relate genes were measured. Compared with the HFD group, AG could significantly decrease the protein and mRNA expression of NF-κB-p65, C/EBPβ and PPARγ in at least the high dose (Fig. 6A, D-F, J-L). To further validate this mechanism, HEPG2 cells processed by ox-LDL were used to verification. Compared with the MOD group, trends in effect of AG treatment approximate with results in animal experiment (Fig. 6B, G-I, M-O). Oil-Red staining is also showed AG dose-dependently attenuates liver lipid deposition in vitro experiment (Fig. 6C, P). This suggests that AG can inhibit the expression of NF-κB-p65, C/EBPβ and PPARγ, and ultimately reduce liver lipid deposition and inflammation.

**Figure6.**
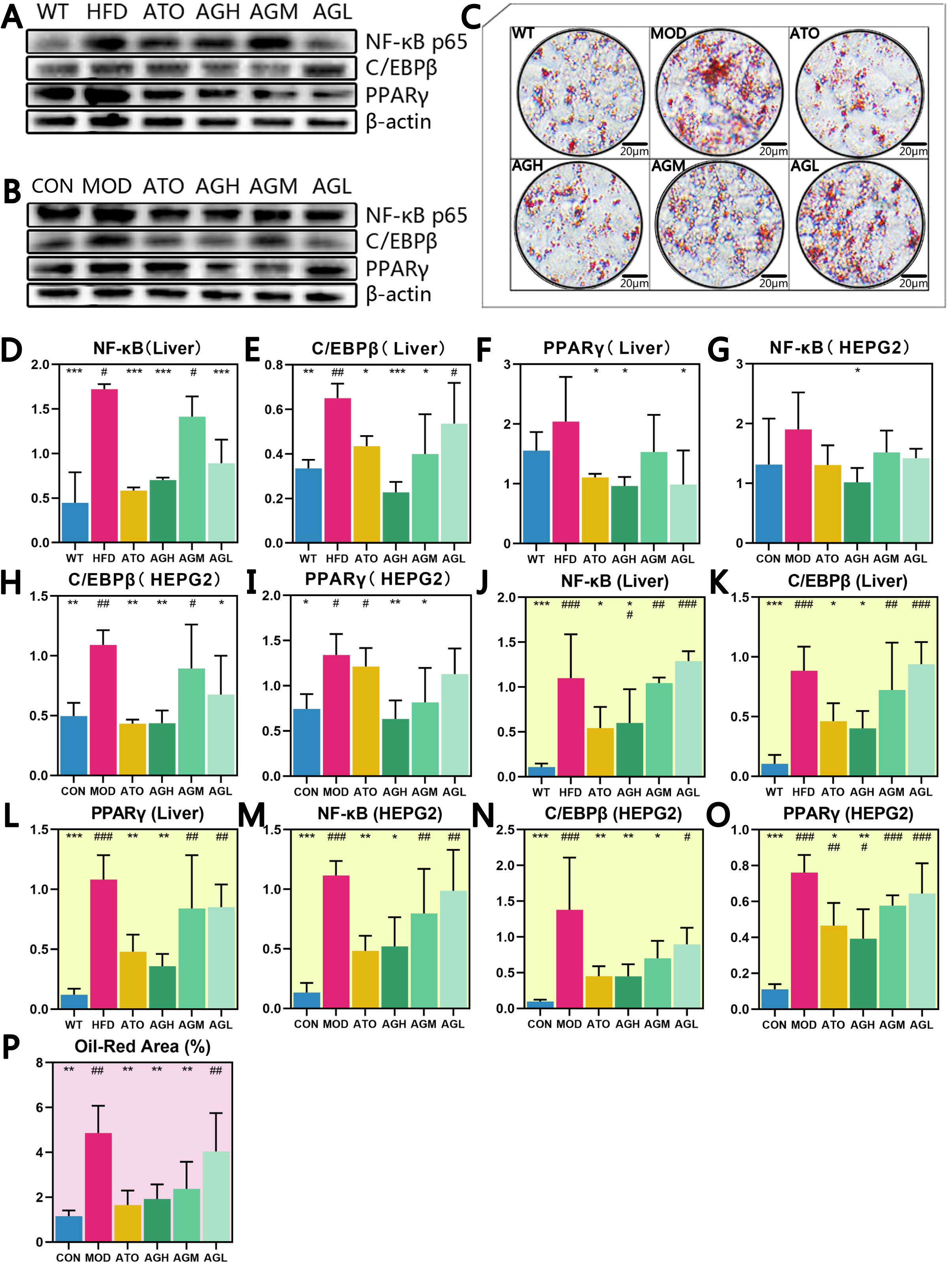
Mechanism of AG attenuates NAFLD. (A) Western blot analyses on liver with anti-NF-κB-p65, anti-C/EBPβ, anti-PPARγ and anti-β actin. (B) Western blot analyses on HEPG2 cells with anti-NF-κB-p65, anti-C/EBPβ, anti-PPARγ and anti-β actin. (C) Oil-Red staining of HEPG2 cells (200× magnification). (D-I) Densitometric analyses of stained membranes, (N=3 per group). (J-O) Relative expression levels of the indicated mRNAs in the liver were determined by qPCR, (N=3 per group). (P) Area of Oil-Red staining (%), (N=3 per group). The data are expressed as the mean±SD of the mean and analysed by One-way ANOVA: *p<0.05, **p<0.01, *** p<0.001 versus HFD group, #p<0.05, ##p<0.01, ### p<0.001 versus WT group.

## Discussion

In our present study, we demonstrated that AG treatment markedly ameliorated HFD-induced ApoE(-/-) mice. In addition to its commonly accepted protective effect on inflammation, we found that the mechanisms through which AG alleviated NAFLD included improve obesity, attenuation of dyslipidemia, partial modulation of intestinal microbiota composition. Furthermore, Liver function in a more prominent security may be one of the advantages it deserves to be selected.

An increasing number of diseases are considered associated with NAFLD. As an underestimated metabolic abnormality, NAFLD is strongly associated with an increased risk of incident prehypertension and hypertension (34). Patients with NAFLD had more than a 2-fold increase in risk of cardiovascular events (CVEs), and individuals with liver fibrosis had a 4-fold increase in risk. In those with NAFLD, liver fibrosis indexes were independently associated with risk of incident CVEs (35). Therefore, it is essential to develop new effective therapeutic approaches to prevent an adverse outcome. AG, a lactones extracted from Andrographis, is known as a small molecular drug against inflammation and fibrosis (36). In the present study, we revealed the effectiveness of AG in alleviating dislipidemia and inflammation, as well as improving obesity, hepatic steatosis and gut microbiota. We observed that ApoE (-/-) mice in C57Bl/6J background fed a HFD developed NAFLD compared with C57Bl/6J mice fed a chow diet, while AG improved the condition. ApoE (-/-) mice on a HFD will be greatly increased blood lipids, which induce liver lipid deposition (37). Research about AG used in the hyperlipidemia model established by ApoE (-/-) mice fed with HFD has not yet been retrieved. Our results demonstrated that lipid metabolism homeostasis was disrupted after HFD administration, and AG restored this metabolic disorder to some extent. The safety and ability of AG to redress the NAFLD in wistar rats on a HFD has also been reported (38). In NAFLD pathway, IL-1, IL-6 and TNF-α are the key inflammatory factors involved in the development of steatohepatitis. Release of mediators were activated by NF-κB, which is one of the key pro-inflammatory signaling pathways in NASH (39). Moreover, inflammatory responses can result in the development of a metabolic disorder. Hence, we tested the levels of inflammatory mediators in different groups. The levels of inflammatory cytokines were increased in HFD group compared with the WT group, while AG administration reduced the expression of inflammatory factors. Collectively, the efficacy of AG on HFD-induced NAFLD involves effects on hepatic steatosis and inflammatory responses.

Regarding the mechanism by which AG attenuates NAFLD, major studies have identically considered that AG functions as an lipid accumulated and inflammatory protectant. Chen et al. found that AG ameliorates lipid accumulation in 3T3-L1 cells by inhibit CCAAT/enhancer-binding protein α (C/EBPα) and C/EBPβ as well as peroxisome proliferator-activated receptor γ (PPARγ). AG derivatives or analogs exhibit potent anti-inflammatory effects in different kind disease models through NF-κB activity (40). Activity of NF-κB, C/EBPβ and PPAR-γ are associated with development of NAFLD (41), (42), (43). It has been reported that C/EBPβ activated PPAR-γ in different model (44), (45). Previous study is also demonstrated that C/EBPβ is an NF-κB-regulated mediator of hepatocellular resistance to TNF toxicity (46). Similarly, we also confirmed that AG could prevent hepatosteatosis induced by the HFD through decreasing NF-κB, C/EBPβ and PPAR-γ in the liver and HEPG2 cells. Cobbina E et al. found that the presence of steatosis, oxidative stress and inflammatory mediators like TNF-α and IL-6 has been implicated in the alterations of nuclear factors in NAFLD (47). Chen HW et al. (48) and Li F et al. (49) found AG inhibit IL-1, IL-6 and TNF-α-induced inflammation in vivo and in vitro disease models. In their study, pro-inflammatory factors was the evaluation index of inflammation -induced disease and the key therapeutic target for AG. Likewise, we also demonstrated that AG could alleviate liver inflammation and serum inflammatory cytokines.

Animal studies in which the gut flora are manipulated, and observational researches in NAFLD patients, have provided considerable evidence that dysbiosis contributes to the pathogenesis of NAFLD (50). Relative research on influence of AG in intestinal flora has not yet found. However, apart from the traditionally reported mechanism, our study primarily revealed that AG could alleviate NAFLD in ApoE(-/-) mice by regulating the gut microbiota. Part of the intestinal flora like Bacteroides participate determines the circulating cholesterol level and may thus represent a novel therapeutic target in the management of dyslipidemia (51). The study by Matziouridou C et al. (52) demonstrated that lingonberries increased cecal relative abundance of Bacteroides in genera level which was down-regulated in HFD-fed mice. We found that abundance of Bacteroides in the HFD group was increased compared with that in the WT group, while AG restored the abundance. Pearson correlation showed it dramatically associated with inflammatory cytokines. Potentially important species such as Faecalibaculum might mediate the beneficial effects of metabolic disturbances in diet-induced obese mice (53). It was reported that Faecalibaculum were identified as the biomarkers for intestinal redox state on hyperlipidemia in HFD-fed mice. AG treatment improved the abundance of Faecalibaculum associated with TG and IL-1α in our study as Fig.5I shown. Moreover, Muribaculaceae abundance was strongly correlated with propionate and community composition was an important predictor of short-chain fatty acids concentrations in mice (54). Cao W et al. found that the intestinal microbiota in HFD-fed mice showed an increased level of Muribaculaceae in emaciated mice compared with fat mice. Unexpectedly in the present study, the AG treated group showed decreased abundance of Muribaculaceae without dose-dependently effect. Mice in ATO group were showed no significant changes in this taxa. Akkermansia is also a kind of beneficial bacteria. In previous studies, Li J et al. (55) reported that daily oral Akkermansia muciniphila attenuates atherosclerotic lesions by ameliorating metabolic endotoxemia-induced inflammation through restoration of the gut barrier in HFD induced ApoE(-/-) mice. Furthermore, Zhu L et al. (56) found berberine treatment significantly reduced atherosclerosis in HFD-fed mice. Akkermansia abundance was markedly increased in HFD-fed mice treated with berberine. Modulation of gut microbiota, specifically an increase in the abundance of Akkermansia, may contribute to the antiatherosclerotic and metabolic protective effects of berberine, In traditional Chinese medicine, Andrographis and berberine have many similar properties especially in tropism of taste. For the first time, our results demonstrated the abundance of Akkermansia was dramatically changed HFD-fed mice and was decreased in AG treated mice. Additionally, Blautia, Alistipes and Ruminococcaceae were reported in previous studies that correlated with HFD-fed models (57) (58) (59), but lack some of the statistically evidence. Despite all this, they were showed a similar trend with the studies. These bacteria might be new candidate phylotypes for predicting and treating metabolic disorders.

## Conclusion

Collectively, our results indicate that AG exerted key beneficial effects on HFD-induced ApoE(-/-) mice NAFLD. In addition, we explored some mechanisms by which the extract might be acting, namely, the modulation of the inflammation involved in metabolic regulation and the alleviation of HFD-induced intestinal dysbiosis alteration. Some intestinal flora, such as Bacteroides, Faecalibaculum and Akkermansia may be fundamentally involved in the mechanism of AG against NAFLD. Our study uncovered possible new mechanisms for AG as a potential therapy for NAFLD. These data provide direction for future studies designed to address the use of AG in clinical settings of NAFLD.

## Abbreviations

NAFLD: Nonalcoholic fatty liver disease;
HCC: Hepatocellular Carcinoma;
T2DM: type 2 diabetes mellitus;
CVD: cardiovascular disease;
CKD: chronic kidney disease;
AG: andrographolide;
NASH: nonalcoholic steatohepatitis;
AS: atherosclerosis;
HFD: high fat diet;
ApoE(-/-): apoE knockout;
TC: total cholesterol;
TG: triglycerides;
HDL-C: high-density lipoprotein cholesterol;
LDL-C: low-density lipoprotein cholesterol;
AST: aspartate aminotransferase;
ALT: alanine aminotransferase;
Cr: creatinine;
BUN: blood urea nitrogen;
IL-1α: interleukin-1α;
IL-1β: interleukin-1β;
IL-6: interleukin-6;
TNFα: tumor necrosis factor α;
H&E: hematoxylin & eosin;
OTUs: Operational Taxonomic Units;
One-way ANOVA: One-way analysis of variance;
PCA: Principal component analysis.

## Authors’ contributions

HYH, CHM, LYD conceived this study and developed the first frame of this manuscript. SS, JXY, and SJJ finish the animal administration and detection. SS drafted the manuscript. SSQ and JQL finish the cell culture and detection. ZQY performed data collection; YGZ and DY revised the manuscript. All authors contributed to the editing of the final manuscript and approved the final version.

## Competing interests

The authors declare that they have no competing interests.

## Availability of data and materials

The data are included in the article as figures, tables, and its additional files.

## Ethics approval and consent to participate

The study was performed in compliance with the animal experimental ethics committee of Guang’anmen Hospital, China Academy of Chinese Medical Sciences (IACUC-GAMH-2019-006). All reasonable efforts were made to minimize animal suffering.

## Acknowledgements/Funding

The Innovative Funding for PhD Students at China Academy of Chinese Medical Sciences (No. CX201904)

